# Neural mechanisms of structural inference: an EEG investigation of linguistic phrase structure categorization

**DOI:** 10.1101/2025.07.01.662085

**Authors:** Qihang Yang, Elliot Murphy, Caimei Yang, Yu Liao, Jianhua Hu

## Abstract

Language comprehension is a complex cognitive process of understanding spoken or written language, during which the brain derives meanings and interprets messages by constructing hierarchical structures from word sequences. A key component underlying this process is structural inference. This computation determines the syntactic category (“head”) of phrases, such as noun phrases (NP) and verb phrases (VP). Categorization confers distinct semantic and distributional properties. However, isolating neural correlates of structural inference is challenging, as modulations of syntactic category usually alter lexical content. Here, we collected the electroencephalography (EEG) data while participants read Mandarin NPs and VPs, which differ in their syntactic “headedness” but with lexical-semantic, lexical-syntactic categories and word order strictly conserved. We found significant theta power increases at left central-parietal scalp sites in NPs relative to VPs around the presentation (0-210 ms) of the nouns, where only the nouns in NPs project a head. Both NPs and VPs relative to one-word phrases exhibited spatiotemporally consistent low-frequency power increases at the point of linguistic composition, consistent with previous studies. Overall, our findings offer novel constraints on neurocomputational accounts of linguistic structure-building through these previously undocumented signatures of syntactic inference.

## 1. Introduction

The capacity to build hierarchical structures (i.e., structure-building) from strings of words is considered a fundamental component in language comprehension (Everaert et al., 2015). The neural basis of structure-building has yielded a number of promising signatures in neurolinguistic studies (Nelson et al., 2017; Zaccarella et al., 2017). Many theoreticians have suggested that (syntactic) structural inference—the ability to infer the categories of a structure, such as noun phrase (NP) and verb phrase (VP)—is crucial in complex recursive structure-building, whereby the identity of constituents is preserved in memory over the course of parsing (Adger, 2012; Chomsky et al., 2019; Hornstein & Pietroski, 2009; Moro & Roberts, 2024; Murphy, 2015). This computation seems to be critical in isolating the brain’s language network, given its domain-specific nature (Berwick et al., 2013). But which neurobiological mechanisms index structural inference? The answer to this question has remained elusive, partly because in real-time processing many distinct syntactic and semantic representations are computed simultaneously (Hagoort, 2019; Pylkkänen, 2019). For instance, the difference between the NP “red boats” and the VP “row boats” exists not only in their different structural categories—a noun phrase headed by a noun, thus an entity; and a verb phrase headed by a verb, therefore an event—but also in their distinct lexical-semantics, lexical-syntactic categories and syntactic structures. Indeed, every time we manipulate lexical content we *also* therefore manipulate syntactic information (Matchin, 2023), since lexical-syntactic category is driving inference. Yet, there remains relatively scarce direct evidence concerning whether the brain computes syntactic processes like structural inference (Boeckx & Theofanopoulou, 2015; Fedorenko et al., 2020; Fedorenko et al., 2012; Murphy, 2024).

Over the past decade, much of the work that informs neurobiological accounts of structure-building relies on comparing structured sentences or minimal-phrases with non-structured stimuli (Bemis & Pylkkänen, 2011; Maran et al., 2022; Nelson et al., 2017; Zaccarella et al., 2017). For one standard paradigm, participants’ brain activations were contrasted when reading sentences (e.g., “the ship sinks”) and wordlists (e.g., “leek mouth ship”). The results of such manipulations have highlighted two left-hemispheric regions for structure-building; the posterior temporal lobe (PTL) and the inferior frontal cortex (IFC). Another paradigm, often known as the “red-boat” paradigm (Pylkkänen, 2020), and typically recruiting electrophysiological measurements, contrasts neural signatures for minimal-phrases and one-word control phrases. This series of experiments found more increased activations for minimal-phrases compared to control phrases in the left anterior-temporal lobe (LATL), starting approximately 200 ms after the onset of the phrasal head (Bemis & Pylkkänen, 2011; Yan et al., 2025). However, follow-up studies suggested the LATL effect is correlated with conceptual-semantic composition (Parrish & Pylkkänen, 2022; Westerlund & Pylkkänen, 2014; Zhang & Pylkkänen, 2018), rather than being a strictly syntactic effect. Nevertheless, recent studies with intracranial recordings using the “red-boat” paradigm also shed light on the role of IFC and PTL, specifically the posterior superior temporal sulcus (pSTS), in minimal phrase composition (Murphy et al., 2024; Murphy et al., 2022), where the IFC is recruited during auditory phrase processing after pSTS and seems more saliently involved in phrasal anticipation and (subsequently) monitoring, rather than generation.

Many current approaches have not been optimized to isolate specific mechanisms of phrasal composition (such as structural inference, nodal depth, or dependency resolution) that underly structure-building. This may be because, in minimal compositional schemes, signatures of lexical-semantic integration may dominate over more rapid processes like phrasal categorization that are also not robustly triggered by task demands (e.g., when the task involves phrase-picture matching). In recent years, studies that focused on the comparison of different syntactic structures have started unraveling some aspects of syntactic processing during structure-building (Flick & Pylkkänen, 2020; Lau & Liao, 2018; Law & Pylkkänen, 2021; Matar et al., 2021; Matchin et al., 2019; Zhao et al., 2025). One common trick used has been to keep differences in lexical-semantics across conditions to a minimum while varying their syntactic structures. For instance, in an innovative study Matchin et al. (2019) highlighted the role of pSTS and the posterior inferior frontal gyrus (IFG) in response to NPs (e.g., “the frightened boy”) and VPs (e.g., “frightened the boy”), where the two phrases share the same words but are distinct in their syntactic structures and word orders. Flick and Pylkkänen (2020) also found distinct activations in PTL to processing noun-adjective pairs in different syntactic relations such as predication (e.g., “Are many trails wide enough …”) and modification (e.g., “There are many trails wide enough…”). The results of such manipulations have unveiled distinct patterns of neural responses to different syntactic structures, contributing to a finer granularity of the mappings between linguistic structure-building and its neural correlates (Poeppel, 2012).

In addition, recent electrophysiology studies have queried the effect of structure-building by focusing on low-frequency neural oscillations (Bastiaansen & Hagoort, 2015; Ding, 2025; Ding et al., 2016; Hardy et al., 2023; Murphy et al., 2024; Segaert et al., 2018; Zhao et al., 2025; J. Zhao et al., 2024). For instance, delta (0.5-4 Hz) entrainment was found to be sensitive to different categorical levels of supra-lexical syntactic structures, such as words, phrases and sentences (Ding, 2025; Ding et al., 2016; Lu et al., 2023). Concurrently, alpha (8-12 Hz) and beta (13-30 Hz) power and phase synchronizations have also been suggested to reflect anticipatory compositional effects (Hardy et al., 2023; Murphy et al., 2022; Segaert et al., 2018), index some form of information pertaining to headed syntactic structure-building (Zhao et al., 2025), and aid syntactic composition via attentional and set-maintenance demands (Hardy et al., 2023; Segaert et al., 2018). For instance, in seminal work Hardy et al. (2023) and Segaert et al. (2018) highlighted the role of alpha power modulations during syntactic composition, when participants listened to morpho-syntactically compositional and non-compositional phrases. While the functional role of theta during structure-building remains an open question, some neurocomputational models of syntax suggest it is related to verbal memory retrieval (Meyer, 2018) and structural inference (Murphy, 2024). One recent neurocomputational model of syntactic structure-building, ‘ROSE’, further invokes mechanisms such as cross-frequency coupling to coordinate spatially disparate computational regimes underwriting compositional syntax-semantics (Murphy, 2024; Murphy, 2025).

The present study follows these lines of research and focuses on the electrophysiological mechanisms of structural inference; that is, how the brain identifies a linear string of words as a hierarchical unit like a noun phrase (NP) or a verb phrase (VP). For instance, the ROSE model provides specific hypotheses about how the brain generate labeled phrase structures, involving theta power modulations being coordinated by low-frequency phase-coupling (Murphy, 2024). Under ROSE, one of the mechanisms through which headedness is established is via theta modulation (via delta-theta phase-amplitude coupling), which leads to a clear hypothesis of theta amplitude modulations differing under varying headedness regimes (Murphy, 2025). Exploiting Mandarin Chinese grammar, we designed a novel paradigm that contrasts the electrophysiological response to NPs and VPs, which differed only in their particles (**Figure 1a**). As such, both phrases in our design were kept the same in their lexical-semantics, word-level categories and word order.

**Figure 1:**
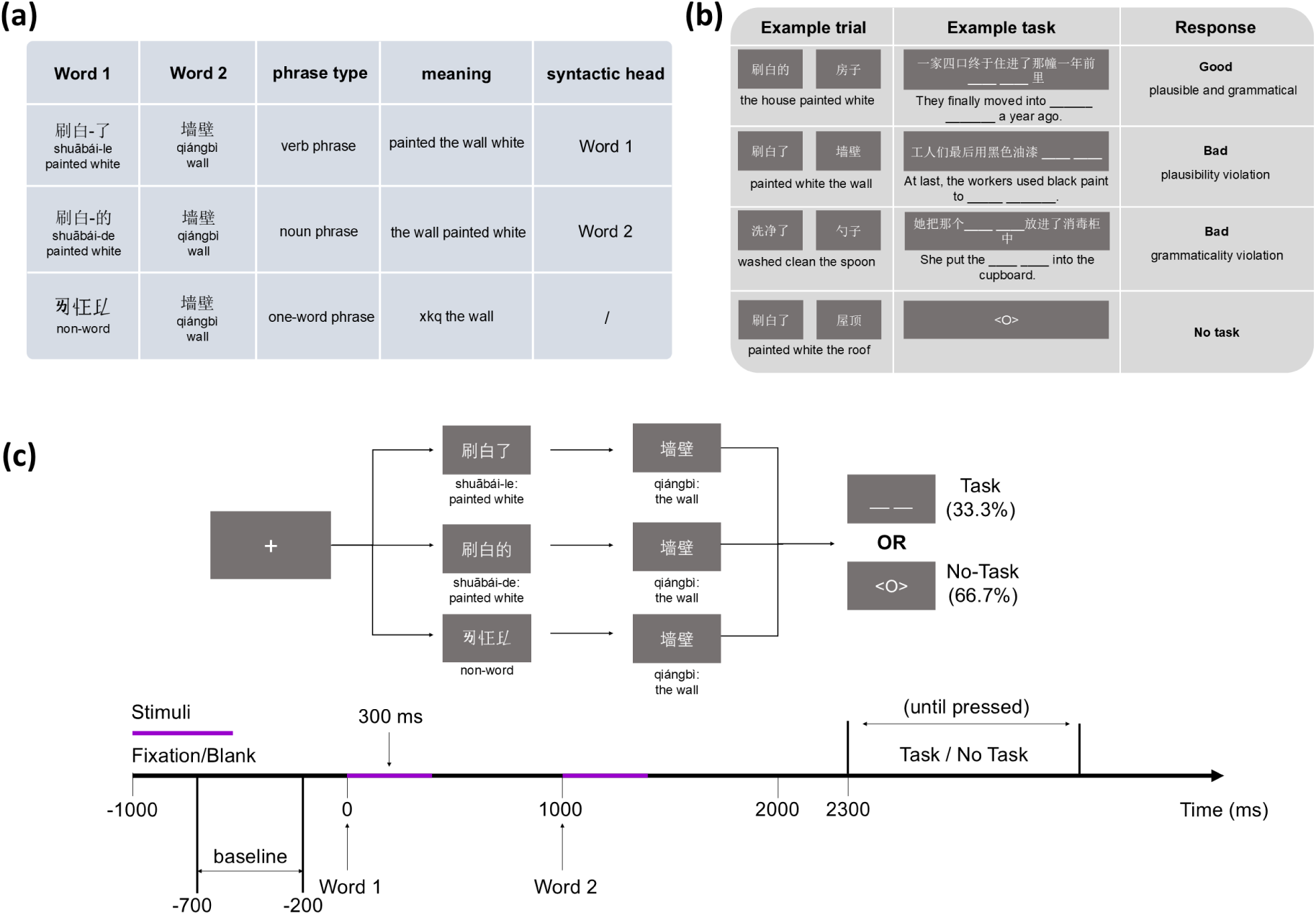
Stimuli design and trial presentation. (**a**) Phrase type manipulations. For simplicity, the first group of characters were labeled “Word 1” and the second group “Word 2”. Word 1 in control conditions consisted of Chinese pseudo-characters, which together constitute a non-word. (**b**) Example task procedure. A third of trials contained task sentences and the remaining two-thirds had a “<O>” displayed at the end of a trial, indicating no-task for the trial. Of the task trials, half had “good” response for the task item (i.e., the preceding stimuli can be filled in the blank to make a plausible and grammatical sentence), and the other half have “bad” responses (i.e., either there is a plausibility violation or grammaticality violation after the insertion of the preceding stimuli). For one-word control phrases, participants were required to judge whether the meaning of the noun can be filled into the sentence. (**c**) Trial structure. The purple line indicates word stimuli presentation time (300 ms) and the black line indicates fixation or blank screen time (300 ms for fixation at the beginning of a trial).

Based on recent syntactic parsing models (Hunter, 2019; Stabler, 2013) and theoretical modeling of structural inference (i.e., properties of the ‘labeling algorithm’) (Chomsky, 2015; Ke, 2023), we expected that the response to the two phrase categories may differ most strongly at the noun, where phrasal head information is most likely to be encoded when composition occurs. Since recent studies have highlighted how neural oscillations can index properties of phrase structure-building, we focused on comparing the time-frequency representations of EEG data in delta (2-3 Hz), theta (4-7 Hz), alpha (8-12 Hz), and beta (15-30 Hz) and gamma (40-80 Hz). We expect to see significant power and/or phase changes in some of these frequency-bands when comparing the response to NPs and VPs, potentially driven by computational regimes driving chunking (delta), categorization or lexical-semantics (theta), attentional modulation during task demands (alpha), anticipatory processing or set-maintenance during workspace updating (beta), and more rapid inferences pertaining to lexical-semantics (gamma) (Kazanina & Tavano, 2023; Meyer, 2018; Murphy, 2024; Prystauka & Lewis, 2019).

In addition, we also included one-word control phrases (“Control”; **Figure 1a**) to more closely model the benefits of the “red-boat” paradigm (Pylkkänen, 2020), in order to compare its response profile to that of NPs and VPs for replicating the well-studied phrasal composition effects. As such, our guiding research questions concerned whether phrasal composition and structural inference can be distinguished by distinct spatiotemporal patterns of oscillatory activity.

Finally, a number of studies have commonly reported event-related potential (ERP) differences between compositional and non-compositional phrases (Fritz & Baggio, 2020, 2022; Neufeld et al., 2016). The amplitude of language-related ERP components like the N400 and P600 increases significantly for compositional phrases relative to non-compositional controls. However, the question of whether these ERP components reflect genuine phrasal composition, or simply task-dependent expectancy or lexical access effects, has remained much debated. We therefore queried this by analyzing the ERPs of the three conditions at the noun (Word 2) where phrasal composition is likely to occur.

## 2. Methods

### 2.1 Participants

Thirty-five right-handed participants were recruited for the experiment. All participants were native speakers of Mandarin Chinese, with no multilingual background and no history of neurological or psychological disorders. Participants received course credit for their participation. Prior to the experiment, written informed consent was obtained from participants.

Data were excluded based on the following criteria: (i) failure to complete all experimental sessions, (ii) for a single condition fewer than 60% of trials remaining after the removal of artifacts (e.g., blinks, eye movements, EMG noise), and (iii) achieving less than 80% accuracy in task response. These resulted in the exclusion of one participant’s dataset for failures to complete the whole experiment, and of five participants’ datasets for containing less than 60% of trials after artifact rejection. The remaining data from 29 participants (8 males, mean = 21 years, SD = 1.57 years, range 18 to 25) were included for further analysis. The experimental protocol was approved by the Ethics Committee of Soochow University, China.

### 2.2 Stimuli

The main experiment consisted of three types of phrases: verb phrases (VP), noun phrases (NP), and one-word phrases (Control). The Control phrases consisted of an unpronounceable non-word with three pseudo-Chinese characters, combined with a noun (see **Figure 1a**). Each phrase in the NP and VP consisted of two Mandarin Chinese character groups. The first group contained three characters, which for reader-friendly concerns we simply labelled as ‘Word 1’, and the second group (Word 2) is a two-character noun.

In the VPs, Word 1 contained a verb and a particle LE. In contrast, in the NPs, it comprises a verb and the particle DE. The particle DE serves as a modifier marker, converting the preceding context into a modifier for the noun; while LE functions purely as an aspect marker for the verb (Feng & Yang, 2019).

Word 2 in all three conditions is a noun, ensuring consistency across conditions. Each verb is used three times in the experiment for both NPs and VPs (each time with different Word 2), resulting in 40 unique sets and yielding 120 items for both conditions. We ensured that all the nouns in Word 2 appeared only once within and across sets to minimize repetition effects.

Prior to the main experiment, a pilot rating study was conducted for all NP and VP items (120 items each) for their naturalness, using a 1-7 Likert scale to measure their comprehensibility. Thirty-one participants who did not participate in the main experiment were recruited. Results from linear mixed-effect models showed no significant difference in naturalness between the two conditions (NP – VP: *beta* = -.007, *t* = -.338, *p* = .738). Following the rating experiment, we re-evaluated the items and excluded two sets that were deemed ambiguous. This resulted in the removal of six items from both the NP and VP conditions, leaving 114 items per condition for the main experiment.

The Control phrases used pseudo-Chinese characters for Word 1 to prevent participants from deriving any meaning or syntactic structure from these stimuli (see **Figure 1a**). The pseudo characters were carefully created by the authors to ensure they do not resemble real Chinese characters or imply any syntactic category. These pseudo-characters were designed to be orthographically plausible but devoid of any semantic or grammatical cues, thus preventing participants from forming expectations based on syntax or meaning. Altogether 38 pseudo character sets were created. Each set of pseudo-characters was repeated three times, resulting in a total of 114 items.

The total number of trials/stimuli was 342, and these were divided into six blocks, with both the block order and the trial order within each block randomized and counterbalanced between participants. Each item appeared only once within a block to avoid repetition suppression effects.

### 2.3 Task and Procedure

Participants were tasked with judging whether the phrase they read within a trial can be filled into a blank of a given “incomplete” sentence frame to form a both grammatical and semantically/pragmatically plausible sentence (Figure 1b). For VP and NP trials, they were instructed to decide if the combination of Word 1 and Word 2 can be filled into the blank slot to form a well-formed sentence. For Control trials, only Word 2 was required to judge the well-formedness of the sentence. One-third of the trials contained this “fill-in-the blank” task, and half of the task items were designed to require a “correct” answer and the other half with an “incorrect” answer. For task items with a “incorrect” response, 52.6% were ungrammatical and 47.4% were implausible sentences when the experimental stimuli were filled into the blanks. In total, 114 task items were constructed, with 57 grammatical ones, 29 ungrammatical ones and 28 implausible ones. For each trial, both response time and accuracy were recorded. Response times were measured from the onset of the task question (for task trials). Accuracy was calculated based on the correctness of the participants’ judgments in the task trials. We counterbalanced the presence of task items and types of resulting sentences, across conditions.

Twenty Mandarin native speakers who did not participate in the main experiment were recruited to norm the task sentences, aiming to verify the appropriateness of the task sentences. A repeated-measures ANOVA on accuracy percentage with Condition (NP, VP, Control) as the within-subject factor revealed no significant effect (*F* (2, 38) = 0.123, *p* = .884; Mean accuracy: NP (94.21%), VP (94.87%), Control (94.73%)).

During the experiment, participants sat in a dimly lit, sound-attenuated room, 70 cm away from the monitor. Stimuli were presented using the E-Prime 3.0 software (Schneider et al., 2002), with a monitor fresh rate of 161 Hz. Participants were told to read carefully the instructions and went through a practice session of 12 trials to familiarize themselves with the procedure and task requirements. They were asked to achieve 80% accuracy in the practice session before proceeding to the recording session.

Participants were visually presented with stimuli, with characters (40-point-font size) in white and the backdrop in grey. At the beginning of a trial, a 40-point-font fixation cross appeared on the screen for 300 ms, followed by a 700 ms blank screen—this represented our baseline period for time-frequency analysis (300 ms to 800 ms, aligned with the onset of the trial). Word 1 was presented on the screen for 300 ms, followed by a blank screen for 700 ms. This manipulation of time interval was chosen based on previous studies to ensure that the lexical processing of Word 1 would likely have been completed within this time window, before the onset of Word 2 (Hardy et al., 2023; Murphy et al., 2022; Poulisse et al., 2020; Segaert et al., 2018). At 2000 ms, Word 2 appeared on the screen for 300 ms and was followed by a blank screen lasting for 1000 ms. After that, one third of the overall trials contained a task sentence displayed on the screen until participants responded. Participants pressed “j” if they judged the sentence would be well-formed and “f” if the sentence was considered ungrammatical or implausible after the fill-in of the stimuli. The remaining two third of the trials had a symbol “<O>” on the screen, indicating no task response, and participants pressed either button to continue. To avoid anticipation effects for the presence of the next trial, a blank screen appeared with a jitter of 500 – 900 ms duration, after participants’ keyboard response. Each participant completed 342 trials (114 NPs, 114 VPs and 114 Control items) in the recording session in a unique randomized order, divided into 6 blocks of 57 trials each. The experiment lasted around 90 minutes.

### 2.4 Data Acquisition and Preprocessing

Raw EEG data were recorded continuously using an actiCHamp (Brain Products, Germany) with a custom-built 64-sensor elastic cap. The sensors on the cap were positioned according to a modified 10–10 system montage (S. Zhao et al., 2024). Two additional sensors, AFz and M1 (left mastoid), served as the ground and online reference sensors during data acquisition, respectively. Horizontal eye movements were monitored using bipolar electrodes placed at the outer canthi of both eyes, while vertical eye movements and blinks were detected with sensors positioned above and below the left eye. The impedances of sensors were kept below 10 kΩ. EEG recording was carried out using the software BrainVision Recorder at a sampling rate of 1,000 Hz.

The continuous EEG data were preprocessed by EEGLAB (Delorme & Makeig, 2004) and Fieldtrip (Oostenveld et al., 2011). Raw data were downsampled to 500 Hz and filtered with a band-pass FIR filter at 0.2-80 Hz. The 50 Hz line noise was identified by visual inspection of channel spectra and removed. The filtered EEG data were re-referenced to the average of the left and right mastoids (M1 and M2) and then segmented into epochs time-locked to Word 2 onset from −2600 ms to 2000 ms. Noisy channels were identified by *pop_clean_rawdata* function and interpolated using spherical spline interpolation (Perrin et al., 1989). An average of 2.6 ± 1.9 (*M* ± *SD*) sensors were interpolated. Then we applied independent component analysis (ICA) on EEG epochs and used *pop_iclabel* function to identify and remove ocular and muscle artifacts (Delorme et al., 2005). An average of 6.6 ± 3.1 (*M* ± *SD*) components were rejected. We applied a baseline-subtraction method using EEGLAB function *pop_rmbase()* to remove channel baseline means from −100 to 0 ms prior to the onset of the fixation cross. Automatic artifact rejection was then performed based on a threshold of ± 75 μV to reject epochs contaminated by residual artifacts. On average, 13.76% of trials were rejected. One-way ANOVA showed no significant differences among NP (mean = 100.2, SD = ± 11.9), VP (mean = 101.1, SD = ± 12.5) and Control (mean = 99.7, SD = ± 12.9) in terms of the number of included trials (*F* (2,84) = 0.088, *p* = 0.916).

After these preprocessing steps, a Laplacian transform was applied to the raw-EEG data using spherical splines (Perrin et al., 1989). The Laplacian is a spatial filter (also known as current source density (CSD) transformation) that serves as a reference-independent normalization method and amplifies localized bioelectric patterns while suppressing diffuse electrical activity. This mathematical approach facilitates cross-study data comparability by converting measurements into standardized units (μV/cm²), thereby reducing methodological variability between research groups (see Carvalhaes and de Barros (2015); Kayser and Tenke (2015a, 2015b) for discussions of the benefits of using the Laplacian spatial filters in EEG data processing).

### 2.5 Statistical analysis

#### 2.5.1 Behavioral data analysis

We used mixed-effects models with *lme4* (Bates et al., 2015) and *lmerTest* (Kuznetsova et al., 2017) in R (version 4.4.3) to examine both response accuracy and reaction time (RT), with accuracy modeled using logistic mixed effects regression and RT using linear mixed effects regression. Reaction times were log-transformed to reduce skewness. Fixed factors include Condition (3-level: NP, VP, Control) and Type (3-level: correct, ungrammatical, unpragmatic) and their interactions. Both factors were sum-coded to facilitate interpretation of main effects and interactions. Subject and Item were selected as random effects, with random intercepts and slopes for Condition and Type for each subject and item. We first excluded 14 trials (< 0.01%) due to extremely short (< 0.2 s) or long RTs (> 20 s). And then for RT analysis we excluded trials whose RTs exceeded ±3 standard deviations from the overall mean RT. Additionally, trials with incorrect responses were removed to ensure only correct-response data were analyzed for RT.

As recommended by Barr et al. (2013), we first included the full random effects structure for each model. When some models with the full random effects structure failed to converge, we simplified them by initially dropping the covariance between random slopes and random intercepts for Items, then for Subjects. If the model still did not converge, we progressively removed the slopes that contributed the least to the variance explained until convergence was achieved.

Models’ assessment was performed with the “performance” R’s package (Lüdecke et al., 2021). Significance of the fixed effects was determined using the Satterthwaite approximation for RT model and Likelihood Ratio Test for accuracy model. Finally, we conducted post-hoc pairwise comparisons with Tukey correction for multiple comparisons. The converged models included Type for RT model and Condition for accuracy model as random slope for Subject: RT ∼ Condition*Type + (1 + Type | Subject) + (1 | Item); accuracy ∼ Condition*Type + (1 + Condition | Subject) + (1 | Item).

#### 2.5.2 Time-frequency power analysis

We analyzed the time-frequency representations (TFRs) of each trial for the 2-80 Hz frequency range in 1 Hz step, deriving a total of 79 frequency bins. Time-frequency decomposition was achieved by convolving a family of Morlet wavelets, with varying number of cycles from 2 to 15 cycles applied to linearly spaced frequency bins. The decomposed data were baseline-corrected to the blank slide appearing after the fixation cross. TFRs of each trial in the three conditions were obtained by subtracting the power in the baseline period (−1700 ms to −1200 ms). TFRs of each condition and participant were calculated separately and then averaged across participants.

In line with previous studies (Bastiaansen & Hagoort, 2015; Hardy et al., 2023; Segaert et al., 2018), the following pre-defined frequency bands were used: delta (2-3 Hz), theta (4-7 Hz), alpha (8-12 Hz), beta (15-30 Hz), and gamma (40-80 Hz). To address the multiple comparison problem, significance was determined using non-parametric temporal-spatial cluster-based Monte Carlo permutation tests (Maris & Oostenveld, 2007). The adjacent sensors regarded as neighbors were determined by using the function *ft_prepare_neighbours* with “triangulation” method in Fieldtrip (Oostenveld et al., 2011). A cluster consisting of at least 3 significant adjacent sensors were deemed as spatial neighbors. We first averaged the power of the frequency bins within its pre-defined frequency band, and then dependent t-tests (two-tailed) were performed on each time point and sensor within a frequency band, and all samples whose t-values were greater than a threshold were selected (sample-level alpha = .05, Bonferroni correction for two-tails). All spatially adjacent sensors with the similar significant time points were grouped into a single cluster. We then performed 100,000 random permutations by randomly shuffling condition labels between the conditions to generate a null distribution, yielding 100,000 clusters. If the largest observed cluster exceeded 95% of the clusters derived from permutations, we concluded that conditions significantly differed within the test window. As there were three pairs of comparisons, namely NP vs. VP, NP vs. Control, VP vs Control, we adjusted the clusters’ p-values via Bonferroni method; that is, the cluster-level alpha was set at 0.017. Hence, we considered clusters with original p-values lower than 0.017 as showing significant differences for a pair of comparison, whereas clusters with original p-values lower than 0.05 but higher than 0.017 as trends towards significance. We first performed the analyses within the time window of interest, centered around the presentation of the second word (−300 to 1300 ms of the epoch). We then extended the analyses to the entire presentation of the stimuli (−1000 to 1300 ms of the epoch, time-locked to Word 2).

To ensure the effects we found in comparing power synchronizations in TFRs across the three conditions were not merely driven by evoked response, the time-frequency representations of ERPs from each participant and condition were derived and performed the same analysis.

#### 2.5.3 Inter-trial phase clustering analysis

Inter-trial phase clustering (ITPC) was obtained by first transforming raw EEG signals into the frequency domain via Morlet wavelets and then computing phase clustering following Cohen (2014):

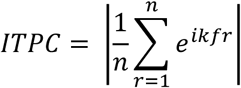

where *n* is the number of trials, *e*^*ik*^ is from Euler’s formula which provides complex polar representation of a phase *k* at a frequency *f* and trial *r* out of *n* trials in total. ITPC is proposed to capture the phase synchrony between individual trials for trial-based design (van Diepen & Mazaheri, 2018). ITPC of each trial for the three conditions across participants was obtained by convolving a family of Morlet wavelets for the 2-80 Hz frequency range in 1 Hz step, with varying number of cycles from 2 to 15 cycles applied to linearly spaced frequency bins. To detect significant time-frequency clusters, a two-tailed cluster-based permutation t-test (sample-level alpha = 0.05) was performed on ITPC difference at the group level, with 100,000 random permutations. Following prior work on syntactic processes and phase synchrony (Brennan & Martin, 2020; Zhao et al., 2025), the test was only run on data averaged over a subset of central sensors (FCz, FC1, FC2, FC3, FC4, Cz, C1, C2, C3, C4, CPz, CP1, CP2, CP3, CP4). We corrected for three comparisons using Bonferroni procedure.

#### 2.5.4 Event-related potentials analysis

Following prior work (Fritz & Baggio, 2020, 2022; Neufeld et al., 2016; Zhao et al., 2025), the event-related potentials (ERPs) were analyzed to find potentially different phrasal composition effects. Specifically, we expected the response to Control to be different from that to NP or VP. We focused on the ERP responses on Word 2. The long-epochs were divided into shorter ones, from −200 ms to 1300 ms, time-locked to the onset of Word 2. Channel baseline means were removed from −200 ms to 0 ms. Each participant’s ERPs were obtained by averaging the epochs separately in the three conditions.

As the question of whether EEG recordings can detect real compositional effects is much debated, and the latency of the claimed “composition effects” in EEG recordings varies within the literature, we conducted non-parametric temporal-spatial cluster-based Monte Carlo permutation tests (two-tailed) for exploratory analysis (Maris & Oostenveld, 2007). The tests were performed on each sensor and time points within the epoch. Parameter settings of the cluster-based statistics for ERPs analysis were the same as for time-frequency analysis, with 100,000 random permutations. Bonferroni method was used for three comparisons.

#### 2.5.5 Effect size analysis

The effect size (Cohen’s *d*) for significant clusters identified by the cluster-based permutation test was calculated by first identifying sensors and time points belonging to each significant cluster. Individual participants’ average power values across these sensors and time points were then extracted. Cohen’s *d* was subsequently computed as the difference between conditions’ mean cluster values divided by the pooled standard deviation across participants.

## 3. Results

### 3.1 Behavioral results

Behavioral data from twenty-nine participants’ behavioral data were included in the analysis after artifact rejection in EEG data preprocessing (see Methods). To evaluate participants’ reaction times (RTs) and response accuracy, we employed linear and logistic mixed-effects regression models respectively. The models included two fixed factors: Condition (3-level: NP, VP, Control) and task sentence Type (3-level: correct, ungrammatical, implausible).

The mixed-effects regression analyses for both RT and accuracy reported significant effects of Condition, Type, and their interactions. Post-hoc analyses indicated the Condition effect was primarily due to significantly shorter RTs and higher accuracy in the Control condition compared to NP and VP (RT: *p* = 2.6·10 ^−10^ and 4.9·10 ^−7^, respectively; accuracy: *p* = .0001 and .0038, respectively). Similarly, the Type effect was driven largely by shorter RTs for unpragmatic sentences, relative to both correct and ungrammatical sentences (*p* = .0014 and 5.8·10 ^−6^, respectively), and lower accuracy in responding to ungrammatical sentences compared to implausible sentences (*p* = .0160).

The significant interaction (**Figure 2**) was explained by notably longer RTs in the NPs compared to both VPs and Control for correct sentences (*p* = 6.6·10 ^−6^ and 1.2 ·10 ^−8^), longer RTs in NP and VP relative to Control for ungrammatical sentences (*p* = 1.7·10 ^−9^ and 1.9·10 ^−7^), lower accuracy in NP to VP and Control for correct sentences (*p* = .009 and .018), and lower accuracy in both NP and VP to Control for ungrammatical sentences (*p* = 2.3·10 ^−5^ and .0005).

**Figure 2:**
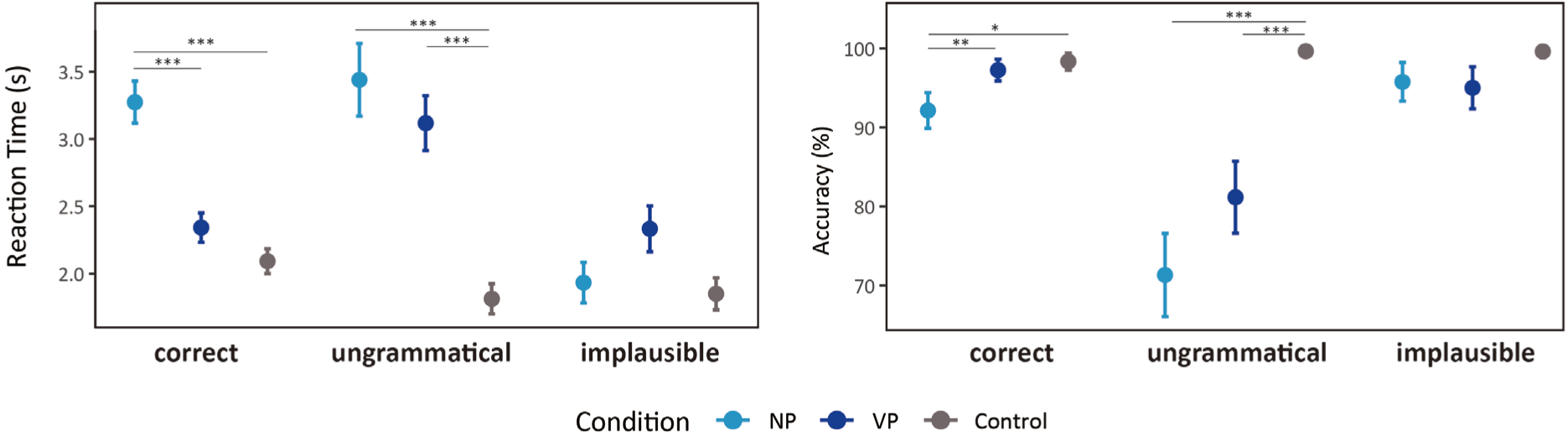
Behavioral results. Mean reaction times (left) and response accuracy (right) across different conditions (NP, VP, control) and sentence types (correct, ungrammatical, implausible). Points represent condition means. Error bars indicate 95% confidence intervals; ***: *p* < .001; **: *p* < .01; *: *p* < .05.

### 3.2 Theta and alpha power modulations for different phrase categories

We first examined whether neural oscillatory activity distinguishes between phrase categories. To this end, we compared TFRs (reflecting event-related power synchronizations) and ITPC (reflecting trial-based phase synchronizations) across pre-defined frequency bands (delta: 2–3 Hz; theta: 4–7 Hz; alpha: 8–12 Hz; beta: 15–30 Hz; gamma: 40-80 Hz).

These comparisons between NPs and VPs were conducted using non-parametric cluster-based permutation tests across all sensors and time points within the analysis window (–300 to 1300 ms), time-locked to the onset of Word 2. We corrected the clusters’ p-values with Bonferroni methods for three comparisons (see Methods). We considered clusters with original p-values lower than 0.017 as showing significant differences for a pair of comparison, and clusters with original p-values lower than 0.05 but higher than 0.017 as trends towards significance.

The most pronounced differences between the two phrase categories emerged in theta power. Specifically, the biggest cluster was identified in an early time window (0 ms to 210 ms) immediately after the presentation of the noun (Word 2), showing enhanced theta-band power for NPs compared to VPs (*p*-adjusted = .018, *t-sum* = 485.17, Cohen’s *d* = 0.71; **Figure 3**). Topographically, this theta effect was predominantly distributed over left centro-parietal scalp regions (**Figure 3b**).

**Figure 3:**
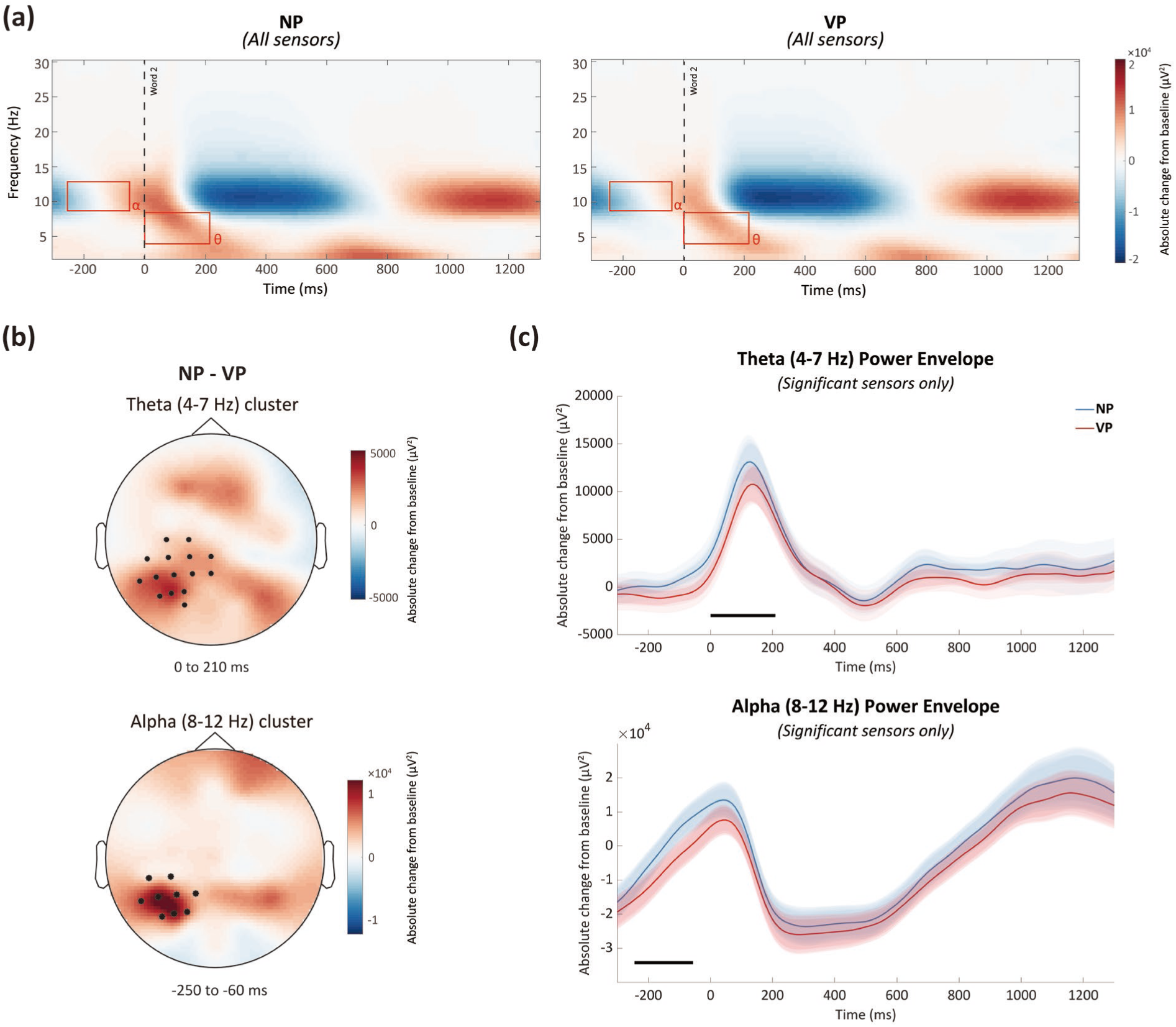
Low frequency power results. Comparison of EEG oscillatory activity during the presentation of noun phrases (NP) and verb phrases (VP). (**a**) Time-frequency representations of power averaged across all sensors, expressed as an absolute change from the baseline period (i.e., −1700 to −1200 ms before the onset of the second word) for the NP trials (left) and VP trials (right). The red rectangles highlight the time period where we observed the significant differences in theta (4-7 Hz) and alpha (8-12 Hz) between the two conditions. (**b**) The scalp topographies of the condition contrast (NP minus VP) of the averaged theta and alpha power activity in two time-windows (0 to 210 ms, and −250 to −60 ms, respectively), where we observed significant power modulations. Sensors with largest effects were highlighted with black dots. (**c**) Theta and alpha power envelop plots. The power envelop displays the time course for the sensors showing significant modulations in power between NP (blue) and VP (red) trials. Significant time periods are highlighted with black lines. The shaded areas in the envelope plots represent ±1 standard error of the mean across participants.

We also observed a trend-level difference in alpha power, with NPs eliciting higher alpha-band power than VPs (*p*-adjusted = .118, *t-sum* = 343.51, Cohen’s *d* = 0.37). Notably, this alpha power modulation occurred prior to the presentation of Word 2 (−250 to −60 ms; **Figure 3b&c**), and shared a similar topographical distribution in the left central-parietal region (**Figure 3b**).

Importantly, the observed theta and alpha oscillatory effects differed from the evoked fields. When we analyzed the TFRs of their ERPs with the same procedure, no significant effects were detected. And no significant differences were observed in the power of delta, beta or gamma frequency bands, both within the time-window of interest and during the whole presentation of stimuli.

Lastly, our ITPC analysis did not reveal any significant differences between NP and VP conditions in any of the examined frequency bands across any time windows.

### 3.3 Oscillatory indices of phrasal composition

To examine oscillatory dynamics specifically associated with phrasal composition, we compared the compositional phrase conditions (NP and VP) with the non-compositional Control condition using the same analysis pipeline, time-locked to the onset of Word 2 and focusing on the main window of interest (–300 to 1300 ms).

Results showed significant low-frequency power increases for both NP and VP conditions compared to the Control condition (**Figure 4a**). Specifically, in the theta-band a significant power increase was observed for VP vs. Control (*p*-adjusted = .048, *t*-sum = 416.69, Cohen’s *d* = 0.83; Latency: 550–730 ms), and for NP vs. Control (*p*-adjusted = .003, *t*-sum = 1384.30, Cohen’s *d* = 0.79; Latency: 460–810 ms). Both effects exhibited a right central-parietal scalp distribution (**Figure 4b**).

**Figure 4:**
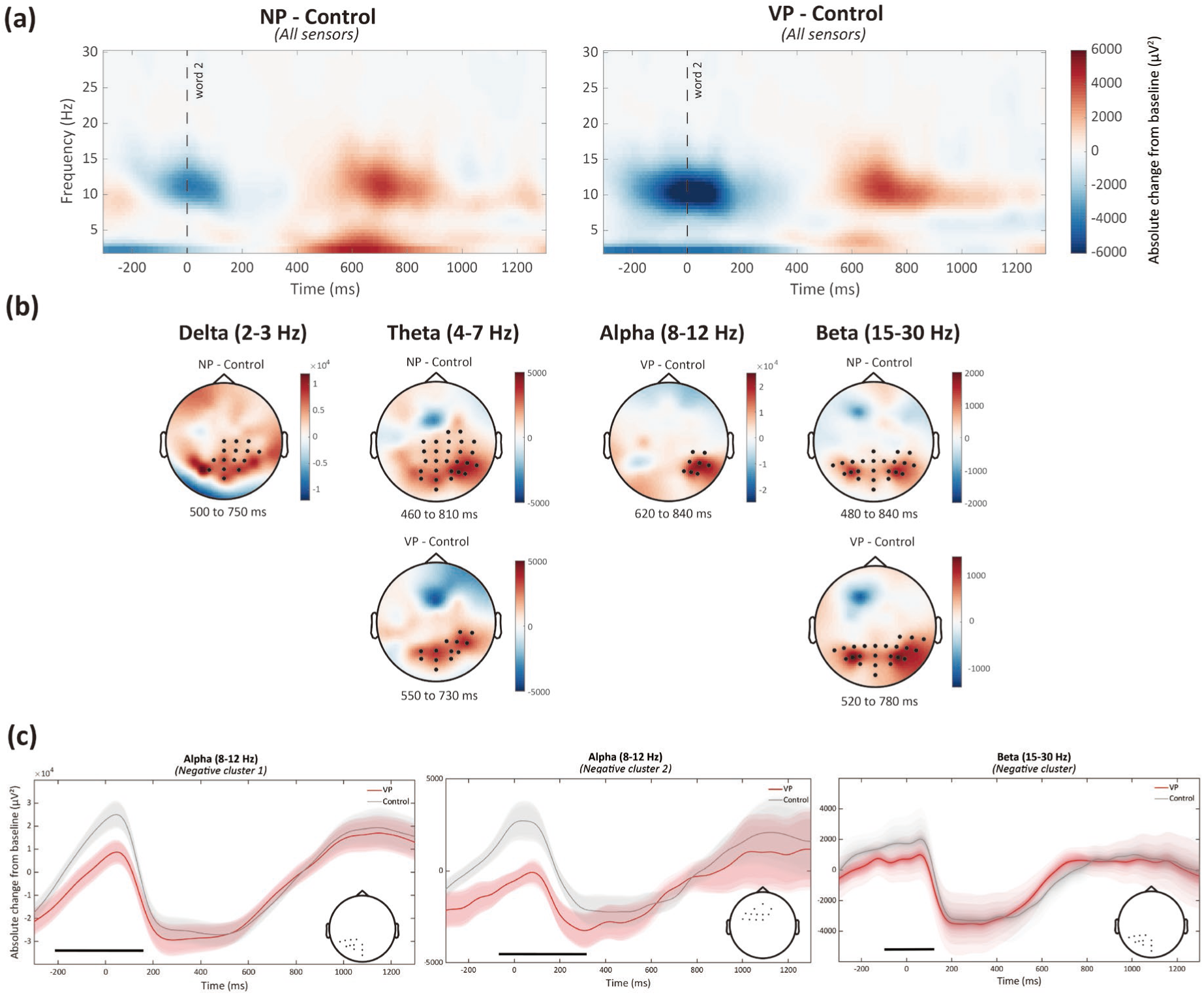
Comparison of the EEG oscillatory activity during the presentation of compositional and non-compositional trials. (**a**) Time-frequency representations illustrating baseline-corrected power differences between compositional and non-compositional phrases, averaged across all sensors. (**b**) Scalp topographies displaying significant oscillatory power modulations across different frequency bands. Sensors with largest effects were highlighted with black dots. (**c**) Power envelopes illustrating unexpected stronger alpha and beta power for the Control trials relative to VP trials, prior to and around the presentation of Word 2. Insets show sensor clusters contributing to these effects. Significant time periods are highlighted with black lines. The shaded areas in the envelope plots represent ±1 standard error of the mean across participants.

In the beta-band, significant power increases were also observed for both VP and NP compared to Control trials, with VP vs. Control showing elevated beta power (*p*-adjusted = .005; *t*-sum = 753.06; Cohen’s *d* = 0.78; latency: 520–780 ms), and NP vs. Control displaying an even stronger beta power modulation (*p*-adjusted = .0006; *t*-sum = 1272.36; Cohen’s *d* = 0.92; latency: 480–840 ms). These beta-band power modulations were bilaterally distributed over central-parietal scalp areas (**Figure 4b**).

Delta and alpha effects were more selective. Specifically, NPs exhibited a significant increase in delta power relative to Control at right central-parietal sensors (*p*-adjusted = .004, t-sum = 1001.57, Cohen’s *d* = 0.92; latency: 500–750 ms), while VPs showed a marginally significant increase in alpha power relative to Control (*p*-adjusted = .096, *t*-sum = 365.59, Cohen’s *d* = 0.59; latency: 620–840 ms), also localized over right central-parietal areas.

Interestingly, significant power modulations emerged in early time windows for the VP-Control comparison: Control trials induced increased alpha/beta power relative to VPs. In the alpha frequency band, two negative clusters survived the test (cluster1: *p*-adjusted = .014, *t*-sum = −808.37, Cohen’s *d* = 0.72, latency: −210–160 ms; cluster2: *p*-adjusted = .023, *t*-sum = −683.65, Cohen’s *d* = 0.75, latency: - 70–320 ms). Topographically, cluster 1 was mainly concentrated over left central-parietal sites, whereas cluster 2 showed a left frontal focus (**Figure 4c**). In beta-band power, a negative cluster was identified (*p*-adjusted = .009, *t*-sum = −620.21, Cohen’s *d* = 0.86 latency: −100 to 160 ms), also predominantly localized over left central-parietal sites (**Figure 4c**). We will address potential implications and interpretations of these unanticipated early power increases for the Control compared to VP stimuli in the Discussion section.

To further explore early compositional processes, we extended the analysis window (–1000 to 1300 ms). In this broader window, consistent power and phase differences between NP vs. Control and VP vs. Control were found during the presentation of Word 1, especially in the delta and theta bands. These early effects were driven by evoked responses.

### 3.4 Event-related potentials

Lastly, we compared ERPs across the three conditions, probing whether phrasal composition effects were sensitive to this metric under task demands distinct from those in the “red-boat” paradigm (Fritz & Baggio, 2020; Neufeld et al., 2016).

Visual inspection of the grand-average ERPs time-locked to the onset of Word 2 indicated that the Control condition elicited larger negative-going waveforms at frontal sites compared to NPs and VPs (**Figure 5a**). Results from cluster-based permutation tests revealed significant differences for both the NP/Control and VP/Control comparisons. Specifically, the NP/Control contrast yielded a positive cluster with a latency of 444–492 ms (*p*-adjusted = .027, *t-sum* = 610.37, Cohen’s *d* = 0.95; **Figure 5b**, top panel). A similar positive cluster emerged slightly later, from 446–524 ms (*p*-adjusted = .009, *t-sum* = 1126.91, Cohen’s *d* = 0.94; **Figure 5b**, bottom panel). Both effects were most prominent over frontal scalp regions, as shown in the topographical distributions (**Figure 5c**). The spatiotemporal distributions of the clusters are consistent with FN400, a more frontally distributed N400 effect (Voss & Federmeier, 2011), which has been associated with lexical-semantic processes.

**Figure 5:**
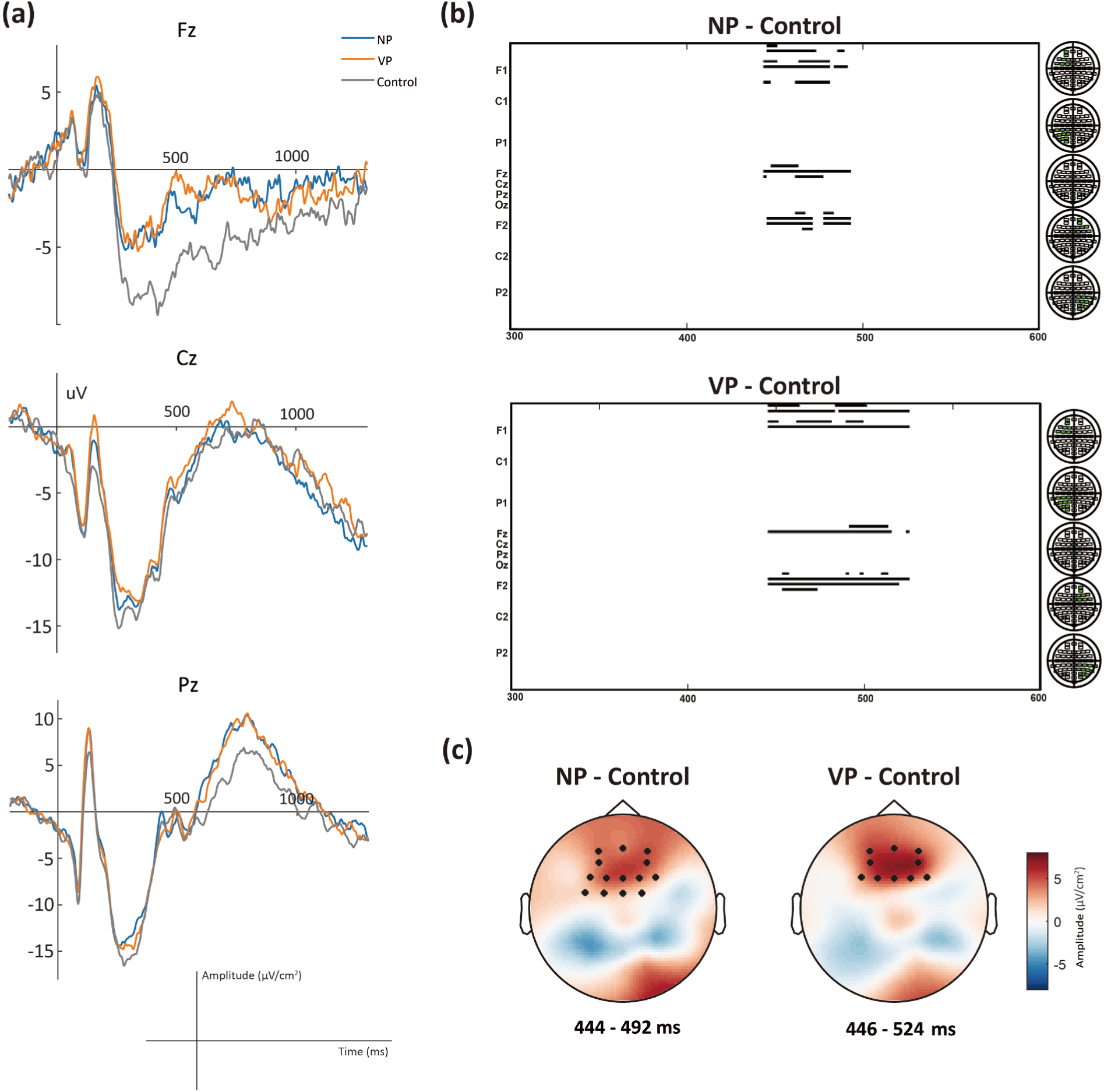
ERPs for NP, VP and Control trials. (**a**) Grand-averaged ERP waveforms time-locked to Word 2 at selected frontal, central and posterior sensors for NP, VP and Control trials. (**b**) Raster plots displaying the spatiotemporal profile of significant effects for NP/Control and VP/Control comparisons. Time periods and sensors showing significant differences for the comparisons are highlighted with black lines. (**c**) Scalp topographies displaying the mean difference (μV/cm^2^) in NP/Control and VP/Control contrasts within the time windows showing significant differences. The black dots illustrate where the effects were largest at the scalp level.

## 4. Discussion

We used a minimally-contrastive reading paradigm exploiting Mandarin Chinese grammar to investigate the neural basis of phrase composition. Phrase composition is a complex cognitive process – involving the establishment of hierarchy, headedness (syntactic category), morpho-syntax, dependencies (agreement, binding), semantic ‘Theta roles’, and lexical-semantic information. Here, we placed specific focus on *headedness*. EEG responses to one-word Control phrases were contrasted with responses to NPs and VPs. Meanwhile, structural inference effects were investigated by directly comparing EEG responses between NPs and VPs, which acutely differ in their categorial features. We found a number of distinct oscillatory signatures marking phrasal composition and structural inference.

First, both NPs and VPs in our design consistently showed increased low-frequency power relative to Control trials in late time windows at ∼ 500 to 800 ms. Specifically, theta and beta power exhibited consistent, significant modulations for NP-Control and VP-Control comparisons. There was also a significant delta power increase in NPs compared to Control, and a trend towards significance in alpha power increase in VPs relative to Control, both in late time-windows and with central-parietal distributions. Delta-band oscillatory activity, specifically its entrainment, has been associated with sequence chunking relevant to typical informational bounds for linguistic phrases (Chen et al., 2025; Ding, 2025; Ding et al., 2016). Theta-band power increase has been suggested to reflect verbal memory retrieval processes in language comprehension tasks (Bastiaansen et al., 2008; Meyer, 2018). Alpha/beta power increases have been found to underpin (peri-)structure-building processes, where structured stimuli (i.e., sentences/phrases) showed more alpha/beta power than unstructured ones (i.e., word lists/one-word phrase), and these bands have also been implicated in syntactic prediction (Bastiaansen & Hagoort, 2015; Bastiaansen et al., 2010; Hardy et al., 2023; Murphy et al., 2024; Segaert et al., 2018). In addition, increasing the structural complexity of a phrase induces more beta power (Matar et al., 2021), highlighting the role of beta-band oscillations in aspects of syntactic processing. In recent neurocomputational models for syntactic composition, beta-band oscillations are suggested to reflect structural maintenance over workspaces (Murphy, 2020, 2024).

Given the design of the present study, the late right central-parietal theta power increase potentially indicates that lexical-semantic or contextual processing (i.e., Word 1) was more demanding for compositional (i.e., NPs and VPs) compared to non-compositional (i.e., Control) phrases. Concurrently, it is possible that the late bilateral central-parietal beta power increases in compositional compared to non-compositional phrases reflect the need for greater cognitive resources for syntactic processes like structure-building and maintenance (Matchin et al., 2024), or the utilization of domain-general cognitive systems for interpretation after structure-building (Nozari & Martin, 2024; Zhang et al., 2025). Altogether, activity modulation in these low-frequency bands potentially underpins various aspects of phrasal composition (from construction, maintenance, categorization, and prediction), consistent with current literature on the neural basis of composition (Hardy et al., 2023; Matar et al., 2021; Murphy et al., 2024; Murphy et al., 2022; Segaert et al., 2018).

Surprisingly, prior to the presentation of Word 2 there was a strong, significant alpha/beta power increase in Control phrases compared to VP trials in left central-parietal (alpha/beta) and left frontal (alpha) sites. There was also a weaker alpha power increase in NPs compared to VPs prior to the onset of Word 2. As mentioned, alpha/beta power increases have been associated with syntactic anticipation prior to composition (Meyer, 2018; Murphy, 2020, 2024), and prior work that focused on minimal-phrase composition also found early left frontal alpha/beta power modulations for compositional stimuli (Hardy et al., 2023; Murphy et al., 2022; Segaert et al., 2018). Thus, it is tempting to interpret the alpha/beta power difference between VP and Control trials as reflecting anticipation of composition. However, the lack of early alpha/beta power modulations in the NP/Control contrast and a trend towards significance in early alpha power modulations in the NP/VP contrast instead suggests that this early effect may reflect the pre-allocation of more general cognitive resources specified for task demands. For instance, rule-switching and set-shifting tasks show coupled alpha/beta power that scales with the number of candidate rules held in working memory (Zioga et al., 2023). Alternatively, phrasal prediction may allocate resources indexed by distinct bands, such that ‘syntactic prediction’ is too generic a process to assign to one fixed canonical band. We leave this topic for future research given that our paradigm was not optimized for addressing these questions of prediction.

As we have indicated, there are some more general framings in the literature that might be relevant for functional interpretations. Fronto-parietal alpha power increases have been proposed to reflect cortical inhibition, functionally related to attentional modulations (Jensen & Mazaheri, 2010; Klimesch, 2012; Klimesch et al., 2007; Van Diepen et al., 2019; Van Dijk et al., 2010), while beta power has been proposed to reflect top-down lexical-semantic and even syntactic predictions (Lewis & Bastiaansen, 2015; Meyer, 2018). It is likely that the more increased alpha/beta power in NP and Control relative to VP prior to the onset of Word 2 reflects different allocation of attentional resources between these conditions. For Control trials, this alpha/beta power increase could be driven by task-related attentional modulations for participants to allocate more resources to retrieve the lexical-semantics of Word 2 (i.e., the noun), given that in the task sentences only Word 2 was needed to complete the sentence. For NP trials, it is likely that Word 2 in NP but not VP plays the role of the syntactic head of the phrase, and therefore requires additional attention to process the word, potentially in the service of parsing operations such as Reduce and tree-structure generation (Hunter, 2019).

In addition, we note that there are low-frequency bands power and phase modulations in NP/Control and VP/Control comparisons, around the presentation of Word 1. We found increased delta, theta, and alpha power and greater phase clustering for Control trials relative to NPs and VPs. These power modulations seem to be driven by evoked response (i.e., the effect is stimulus-driven). Given that Word 1 is an unpronounceable non-word in Control phrases, this increased low-frequency power may indicate greater processing effort for one-word Control phrases, i.e., greater lexical-phonological effort at Word 1 for control trials (potentially due to attempted lexical access or heightened phonological compute), or it may simply indicate an effect of visual repetition suppression (Murphy et al., 2022). Inter-trial phase clustering has also been a reliable signature of lexicality classification (Jensen et al., 2019). Together, in this design, modulations in power and phase clustering around Word 1 indicate electrophysiological signatures underpinning non-word and word processing.

Taken together, within the context of our paradigm we suggest that low-frequency power modulations in late time windows after the presentation of Word 2 reflect some part of genuine phrasal composition effects, while early alpha/beta oscillatory responses index attentional modulations by task demands, rather than strictly indexing syntactic anticipation of composition.

Another component of our results concerns the early left central-parietal theta power modulations between NPs and VPs. We first exclude the possibility that this effect reflects verbal memory retrieval (Meyer, 2018) and/or properties of different syntactic structure processing. Indeed, in our design, NPs and VPs differ not only in their syntactic heads, but also fundamentally in their syntactic structures. In theoretical linguistics, NPs with pre-nominal modifiers are adjunction structures, whereas VPs are head-complement structures, plausibly requiring more salient syntactic labeling than adjunction structures, which involve Adjoin-type operations (Hornstein, 2008). For instance, Matchin et al. (2019) reported increased activity to VPs (e.g., “frightened the boy”) relative to NPs (e.g., “the frightened boy”) in pSTS and posterior IFG, which they interpreted as reflecting different syntactic processing of head-complement and adjunction structures. Thus, it is tempting to attribute the early theta power modulations to different syntactic structure processing. However, we note that there were no similar early theta power effects in the VP/Control or NP/Control comparisons. Our Control phrase contained only one real word, and if this effect reflected verbal memory retrieval or syntactic structure processing, both NPs and VPs should have shown similar power modulations when compared to Control. The subtlety of processing different syntactic structures is potentially reflected by participants’ behavioral performance: NPs relative to VPs exhibited longer reaction times and lower accuracy in the “correct” task sentences, suggesting greater processing effort in adjunction structures (i.e., NPs).

Theta power modulations have also been associated with selective attention and cognitive control. One study (Klimesch et al., 1997) found increased frontal-central theta power synchronization when participants monitored auditory and visual sequences simultaneously, suggesting that the frontal-central theta activity could be a marker of multisensory divided attention. Another study (Cavanagh & Frank, 2014) proposed that frontal-central theta activity profiles “appear to reflect a common computation used for realizing the need for cognitive control”. Given that the theta power modulations we report had a left central-parietal distribution for NP/VP comparisons, this may be driven by distinct processes from attention or cognitive control, which usually have a more frontal focus.

Following recent neurocomputational models of language comprehension (Kazanina & Tavano, 2023; Martin, 2020; Meyer, 2018; Murphy, 2024), we suggest that the increased early theta power in left central-parietal sites encodes syntactic head information, which is an essential component of structural inferences. For instance, the ROSE model proposes that delta-theta phase amplitude coupling indexes recursive supra-lexical structural inferences, with the lexical-level theta-complex that determines headedness showing greater phase-amplitude coupling with delta phase compared to theta-complexes that are non-heads (e.g., phrasal complements), thus leading to a prediction of theta-power modulation differences between heads and non-heads (Murphy, 2024). Other literature has emphasized the role of delta activity in rule-based hierarchical structure-building (Ding, 2025; Ding et al., 2016; Lu et al., 2023). One study (Zhao et al., 2025) found increased delta/beta inter-trial phase coherence when the word marked the boundary of a phrase, indicating that delta/beta phase is sensitive to phrase closure. This may explain why our ITPC analysis failed to find significant differences in low frequency phase synchronization between NPs and VPs, since Word 2 in both phrases marked the phrase boundary. Regarding the functional role of theta activity in structural inference, the ROSE model suggests that theta-driven dynamics constitute a ‘hand-off’ of information after lexicality has been established, transitioning from encoding lexical memory to a more complexed multi-object syntactic workspace represented in traveling delta and theta waves (Murphy, 2024). In other words, theta power as in delta-theta phase-amplitude coupling may encode the memory of the syntactic head of a multi-word phrase. Given our design, Word 2 acts as the syntactic head in NPs, and as complement in VPs, and an increase of theta power was found in NPs relative to VPs.

Finally, we found an increased FN400 (a frontally-focused N400 component) for Control phrases relative to NPs and VPs. The FN400 is an ERP component typically elicited in recognition memory paradigms. Traditionally, its amplitude is reduced for correctly recognized items compared to new items—a pattern long interpreted as reflecting the operation of familiarity-based recognition (Guo et al., 2013; Stróżak et al., 2016). In addition, there is also evidence that the functional role of the FN400 and N400 is the same (Li & Guo, 2020; Voss & Federmeier, 2011). In general, N400 reflects the difficulty of semantic integration or the pre-activation of lexical-semantic information (Lau et al., 2008). The reduced amplitude of FN400 in NPs and VPs relative to Control trials indicates that access to the lexical-semantics of Word 2 was easier for compositional than non-compositional phrases. It is likely that the verbs in Word 1 in NPs and VPs have selection constraints on Word 2. For instance, the verb “paint” is more likely to be combined with concrete nouns like “cars” and “walls”, than with abstract words like “dreams” and “ideas”. As is typical, verbs will constrain the selection of nouns, narrowing down the types of nouns licensed for composition, thereby facilitating access to the lexical-semantics of Word 2. This suggests that the FN400 effect in the present study indicates lexical-semantic processing rather than genuine composition, an interpretation consistent with prior studies in which phrasal composition may not be indexed by ERP components (Fritz & Baggio, 2022).

In conclusion, this work unveiled novel signatures of syntactic phrase composition centered on the parsing of categorial headedness. This is a key representational ingredient that the language faculty contributes to higher-order thought and combinatorics, and an essential feature for delivering more complex inferences to general semantic systems. As such, future studies should continue to isolate spatiotemporal dynamics of structural inferences via headedness, ideally by expanding on our power-based dynamics and investigating properties of inter-areal dynamics, cross-frequency coupling, and other related measures.

## Acknowledgements

This research was supported by the National Social Science Fund of China under Grant 23BYY170 and under Grant 23&ZD319. We would like to thank: Chenxi Fu, Xiaoyi Wang, Ruyi Yang, Chunlei Zhang, Ran Xu, Qiqi Wu, Xuan Shi for help with data collection, and Ruizhe Zhou for helpful discussions.

